# Land-use change from food to energy: meta-analysis unravels effects of bioenergy on biodiversity and amenity

**DOI:** 10.1101/2021.06.16.448590

**Authors:** Caspar Donnison, Robert A. Holland, Zoe M. Harris, Felix Eigenbrod, Gail Taylor

**Affiliations:** Department of Plant Sciences, University of California, Davis, 95616, California USA; School of Biological Sciences, University of Southampton, Southampton, SO19 1BN, UK; School of Geography and Environmental Science, University of Southampton, Southampton, SO19 1BN, UK; Centre for Environment and Sustainability, University of Surrey, Surrey, GU2 7XH UK

**Keywords:** bioenergy crops, biodiversity, ecosystem services, visual amenity, meta-analysis, systematic review, BECCS, social license to operate, landscape

## Abstract

Most decarbonization scenarios of energy systems necessitate more than 500 Mha of land converted to non-food bioenergy crops to provide both energy substitutes for fossil fuels and negative emissions through bioenergy with carbon capture and storage (BECCS). Understanding the environmental and societal impact of this significant land-use change (LUC) is important in determining where and how bioenergy crops should be deployed, and the trade-offs and co-benefits to the environment and society. Here, we use two systematic reviews and a meta-analysis to assess the existing literature on impacts that are likely to have an important effect on public perceptions of the acceptability of such land use change: biodiversity and amenity value. We focus on the impact of LUC to non-food bioenergy crops on agricultural landscapes, where large-scale bioenergy planting may be required. Our meta-analysis finds strong benefits for biodiversity overall (up 75 % ± 13 %), with particular benefits for bird abundance (+ 81 % ± 32 %), bird species richness (+ 100 % ± 31 %), arthropod abundance (+ 52 % ± 36 %), microbial biomass (+ 77 % ± 24 %), and plant species richness (+ 25 % ± 22 %), when land moves out of either arable crops or grassland to bioenergy production. Conversions from arable land to energy trees led to particularly strong benefits, providing an insight into how future LUC to bioenergy crops could support biodiversity. There were inadequate data to complete a meta-analysis on the effects of bioenergy crops on landscape amenity value, and few generalizable conclusions from a systematic review of the literature, however, findings highlight the importance of landscape context and planting strategies in determining amenity values. Our findings demonstrate improved farm-scale biodiversity on agricultural land with bioenergy crops, but also limited knowledge concerning public response to this land use change which could prove crucial to the effective deployment of bioenergy crops for BECCS.

## Main

Bioenergy has been identified as a key contributor to future energy scenarios consistent with the Paris Agreement targets, and is relied upon in scenarios both with and without bioenergy with carbon capture and storage (BECCS), owing to the multiple ways in which bioenergy can subsitute fossil fuels^1^. It has been estimated that to meet the future demand for bioenergy, large-scale deployment of non-food bioenergy crops, usually fast-growing trees and grasses, will require several hundred million hectares (Mha) of land, and much of this may come from conversion from agricultural uses, including arable land currently used for food and feed crops and grassland for animal production^2,3^. To meet a 2 °C target land-use estimates for BECCS alone stand at 380-700 Mha^4^, equivalent to a land area 1-2 times the size of India^5^.

This scale of land-use change (LUC) to bioenergy crops is a potential threat to water and food security, sustainable development, and biodiversity, although co-benefits may also exist^6^. There is evidence that conversion from agricultural land-use to non-food bioenergy crops can both enhance and degrade environmental processess and services^7–10^, but much of the prior empirical research has focused on regulating services such as changes in soil carbon and greenhouse gas (GHG) mitigation potential^11,12,13^. Potentially important impacts of bioenergy crops on the wider environment, which could prove critical to the success of future bioenergy policies, are difficult to measure and are therefore poorly represented in land-use and environmental impact modelling tools^14,15^.

LUC represents the major driver of biodiversity loss in recent decades, with 47 % of natural ecosystems in decline and around one million plant and animal species estimated to be at risk of extinction^16^. Understanding the impact of expanded bioenergy cropping on different species is crucial and previous qualitative reviews have identified both positive and negative impacts of bioenergy crop deployment on biodiversity, depending on the reference land-use and land management^7,17–20^. The conversion of natural and semi-natural ecosystems to bioenergy crops is likely to lead to negative impacts on biodiversity^19,20^, as well as the loss of stored carbon^12,13^. It is often assumed that an expansion of bioenergy crops will largely occur on marginal unimproved semi-natural land rather than productive agricultural land, but this assumption is likely too simplistic. Whilst sizeable opportunities for bioenergy crop deployment may exist on land that does not conflict with either natural ecosystems or food production^21^, in reality, these areas of land may not align with locations of bioenergy demand, adding to both financial and carbon costs^15^. These areas may also be difficult for crop cultivation or prioritised for future food security over conversion to bioenergy cropping^22^. With farmland used for crop and livestock production currently covering one third of the land surface of the earth^16^, we posit that if natural ecosystems are to be protected, agricultural land will be required for at least some of the increased future bioenergy cropping and that this could support biodiversity and help reverse recent trends in species loss. Whilst many studies point to an inevitable conflict between land for food and for bioenergy^23^, it is also possible that agricultural land will become available through increased precision agriculture and higher crop yields, alongside reductions in food losses and changes in dietary trends and associated declines in livestock and livestock feed requirements^24,25^.

Societal impacts of bioenergy could determine the success or failure of technologies such as BECCS. Policy discussions of BECCS are taking place whilst public understanding and public debate of the technology remains limited^26,27^. This presents a clear risk: if BECCS proves politicaly unacceptable then policymakers will need to find alternative solutions to meet the Paris targets in the short time period remaining. Existing BECCS research has provided very limited understanding of impacts at the national and local scale, despite this being exactly where decision-making and public attitudes may determine whether or not the technology is pursued^15,28^. Onshore wind energy, a more mature technology than bioenergy, has faced challenges owing to landscape impact^29^ and there are similar concerns in some communities that bioenergy crops may have a negative impact on the visual landscape^30–32^. The local context of a proposed energy project, engagement with the local community, and communication of the costs and benefits of the project to that community are all considered essential pillars of achieving a Social License to Operate (SLO): the ongoing community support of a technology or activity^33^. There is a need to assess all the evidence for the landscape impact of bioenergy crops to determine whether future bioenergy technologies can achieve public support and a SLO. As well as impacting biodiversity objectives, the effects of bioenergy crops on biodiversity may also be key to the success or failure of a SLO; however, no meta-analysis has yet been applied to the conversion of agricultural land to bioenergy crops in relation to biodiversity. In addition, no systematic review of the visual and recreational impacts, together termed amenity impacts, of land-use change to non-food bioenergy cropping exists. We address both gaps in the literature using a meta-analysis of the impacts of agricultural land-use change to non-food bioenergy cropping on biodiversity and a systematic review of the same land-use change on amenity impacts.

## Materials and Methods

### Systematic Review

Our two systematic reviews followed established protocols^34^. In the biodiversity systematic review, studies were assessed on the inclusion of the following: 1) primary data of the biodiversity impact of agricultural land-use (arable or managed grazing, defined to include permanent, semi-permanent, and rotation grassland) change to bioenergy crops; 2) assessment of a non-food bioenergy crop (*Miscanthus*, *Panicum viegatum* (switchgrass), other energy grasses, short-rotation poplar and willow energy trees; 3) provision of response data for both the treatment (bioenergy) and control (agricultural land). We also limited studies to temperate regions (excluding the polar circles and subtropic regions), exluding tropical crops and regions as outside the scope of this current study. For the amenity and recreation systematic review we set the following inclusion criteria: 1) primary data of the visual or recreational impact of LUC to bioenergy crops; 2) assessment of non-food bioenergy crops as defined above; and 3) a location in a temperate region.

Search strings were first tested for their success in yielding papers identified as relevant to the study. The final search strings were used in the Web of Science and Scopus search engines (see Table S1). Our systematic approach to the peer-review literature was augmented with targeted search of the ‘grey’ literature, using Google Scholar and visiting websites of relevant organisations. The results were downloaded into Excel spreadsheets where title and abstract reviews were completed, with the removal of papers failing to meet inclusion criteria (see Figure 1).

**Figure 1.**
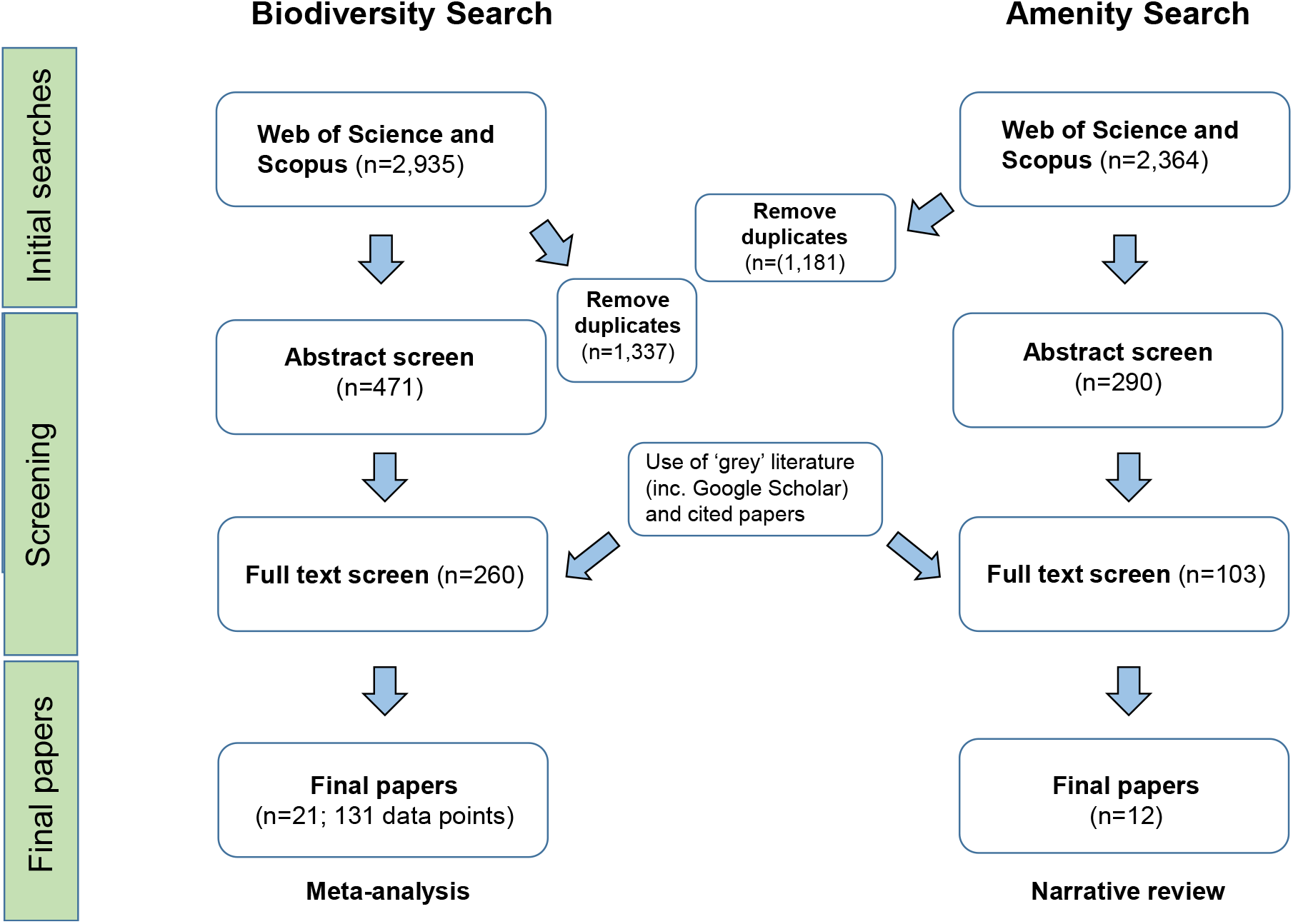
Flowchart of the review process of the two separate systematic searches of the biodiversity and amenity impacts of bioenergy crops. Both searches were conducted in July 2020.

The biodiversity search of Web of Science and Scopus (conducted in July 2020) yielded 2,679 results, in additon to grey literature searches, and 21 papers were found suitable for meta-analysis after title, abstract, and full paper review process. The studies in these papers included data collected on a range of species and we split these based on their taxonmnic coverage between four groups: birds, plants, arthropods, and below-soil organisms (including earthworms and microbial biomass). The studies used typically compared several bioenergy crops with at least one control and measured the impact on more than one species group. For each combination of bioenergy crop, control land-use, and species group, an entry was made into the Excel data table. The 21 papers resulted in 131 observations of relevant data for the meta-analysis. We collected data on the sample size (n), standard error (SE), and mean (see SI). Where data were displayed in graphical form only, the software Data Thief^35^ was used to extract numerical values. We contacted authors where relevant data were missing.

The amenity systematic review (conducted in July 2020) yielded 2,064 results from Web of Science and Scopus, in addition to grey literature searches. This was reduced to 12 papers after the title, abstract, and full paper review process, with very few papers addressing landscape amenity and many of those that did proving unsuitable because they did not study bioenergy crops. Further quantitative analysis was not possible owing to a paucity of quantitative data so a narrative review of the final papers conducted in order to draw out relevant themes and conclusions.

### Meta-analysis

Using the software OpenMEE^36^ we ran one ‘global’ meta-analysis for all species groups combined in addition to separate meta-analyses for each species group and individual species where data were sufficient, for both abundance and species richness metrics. This led to nine separate analyses: all biodiversity (n=104), bird abundance (n=38), bird species richness (the number of distinct species observed; n=19), *Alauda arvensis* (Eurasian skylark) abundance (n=22), *Emberiza* genus (buntings) abundance (n=5), plant species richness (n=8), arthropod abundance (n=17), earthworm abundance (n=5), and microbial biomass (n=17). In this study, the effect size represented the biodiversity change in the treatment (bioenergy) group compared to the reference land-use (agricultural land-use) group. A log response ratio was calculated to represent the effect size: the natural log of the ratio of the mean value of the treatment (bioenergy) to the mean value of the control (arable or grassland). The log response was considered a more appropriate response metric than calculating the effect size using the standardised difference between group means (e.g Hedges’ g) because it does not use within-group variance in its calculation^37^. This is important since variance can vary notably between the studies owing to differences in study design such as geographic location, distribution, and taxonomic group^37^.

The studies were weighted according to the inverse of individual study variance, and thus greater weight was given to larger studies with more precise effect estimates. A grand mean of all the log response effect sizes was calculated using a random-effects model, with the assumption that the true effect size varies between studies and that there is not one single true effect size (when a fixed-effects model is used). We acknowledge that some of variation between results was the result of study heterogeneity, including field size, time of year, and sampling method. However, datasets including this information were incomplete and not included in our analysis. We ensured that between-study heterogeneity was significant using the Q statistic (see SI for details). We also tested for publication bias - a bias towards the publication of positive results - using the funnel plot ‘trim and fill’ method^38^, and asessed studies for evidence of pseudoreplication (see SI).

### Landuse change to bioenergy: impacts on biodiversity

Overall, we found that LUC from cropland and grassland to bioenergy cropping had positive effects on biodiversity, with species abundance increasing 73 % (± 17 %) and species richness rising 80 % ± 24 %, when assessing all studies (Figure 2). Bird abundance was increased by 81 % ± 32 % (n=38, Figure 2) in bioenergy cropping landscapes compared to agricultural land-use, either arable or grassland. Bird species richness also increased, by 100 % ± 31 % (n=19, Figure 2). Soil microbial biomass (n=17) increased under LUC to bioenergy cropping, by 77 % ± 24 % (Figure 2). Arthropod abundance (n=17) was 52 % ± 36 % greater under bioenergy crops, and plant species richness also increased under bioenergy planting compared to arable and grassland cropping, 25 % ± 22 % greater (n=8; Figure 2). Whilst meta-analysis results for earthworm, *Alauda arvensis* (Eurasian skylark; Figure S9), and *Emberiza* (buntings; Figure S10) were not sigificant, a number of the studies reviewed found positive impacts of LUC to bioenergy crops (37 % ± 60 %, p=0.19; 18 % ± 61 %, p=0.50; and 158 % ± 271 %, p=0.16, respectively). Detailed analysis was completed to elucidate the biodiversity impact of specific land-use changes: we found a greater observed increase in biodiversity under conversion to short-rotation energy trees compared to energy grasses, with particularly notable benefits for birds under conversions to energy trees (Figure 3), although conversions to energy grasses were not statistically significant for birds. Bird biodiversity was also supported more under conversions from arable land compared to those from grassland (Figure 3).

**Figure 2.**
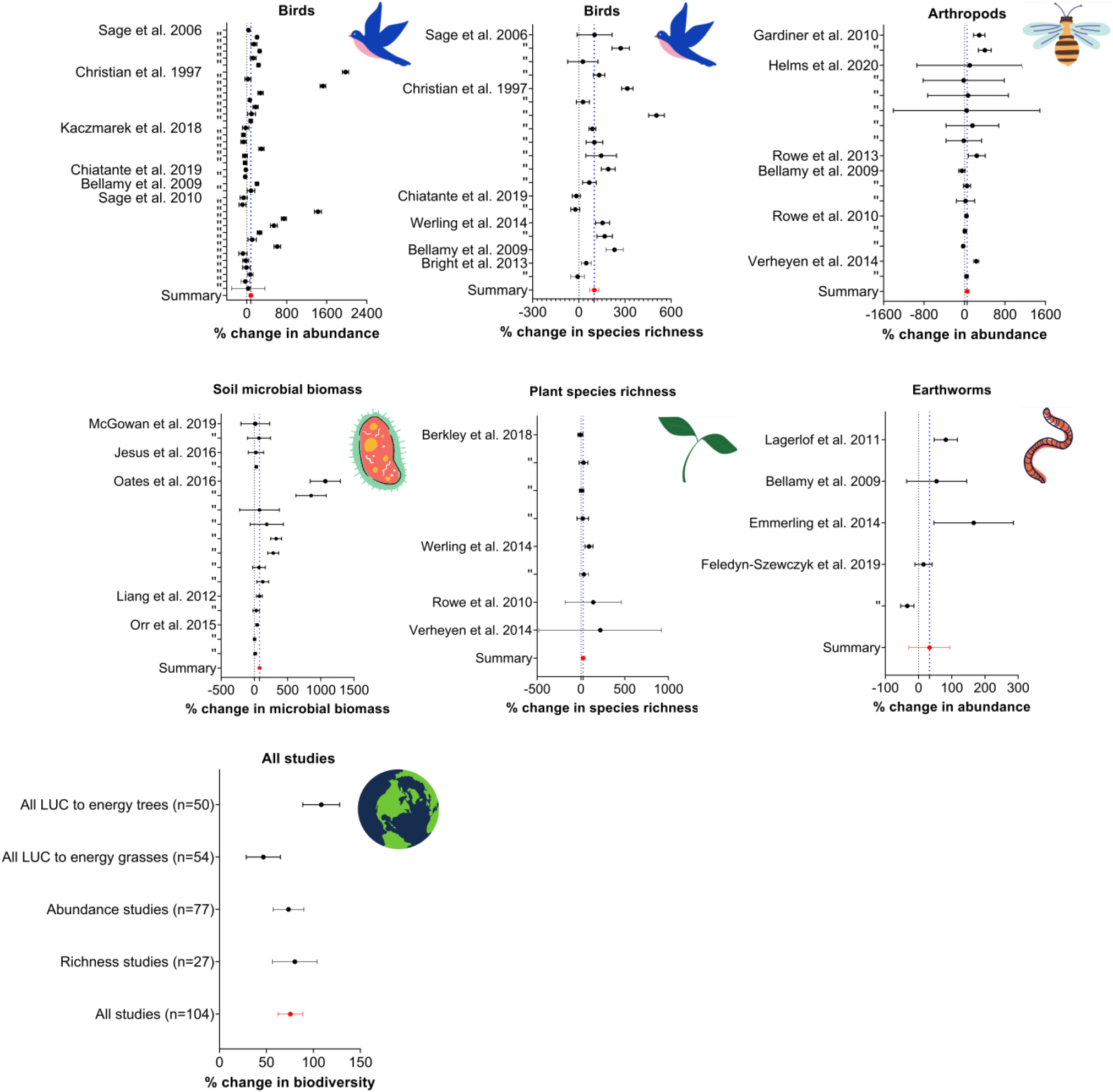
Meta-analyses of the impact on biodiversity taxonomic groups (bird abundance, bird species richness, arthropod abundance, microbial biomass, earthworm abundance, and plant species ricness) of LUC from agricultural land (arable and managed grassland) to non-food bioenergy crops (*Miscanthus*, switchgrass, prairie grass, short-rotation energy trees poplar and willow). Black circles represent mean values of individual study results, with 95 % CI, and red circles represent summary values, with 95% CI. Green dotted line shows overall summary value (weighted average) of each meta-analysis. Bird abundance increases by an average 81 % (± 32 %), bird species richness rose an average 100 % (± 31 %), insect abundance increased an average 52 % (± 36 %), soil microbial biomass increased an average 77 % (± 24 %), and plant species richness increased 25 % ± 22 %. All results in Figure 2 were statisitically significant with the exception of results for earthworms (+ 37 % ± 60 %, p=0.19).

**Figure 3.**
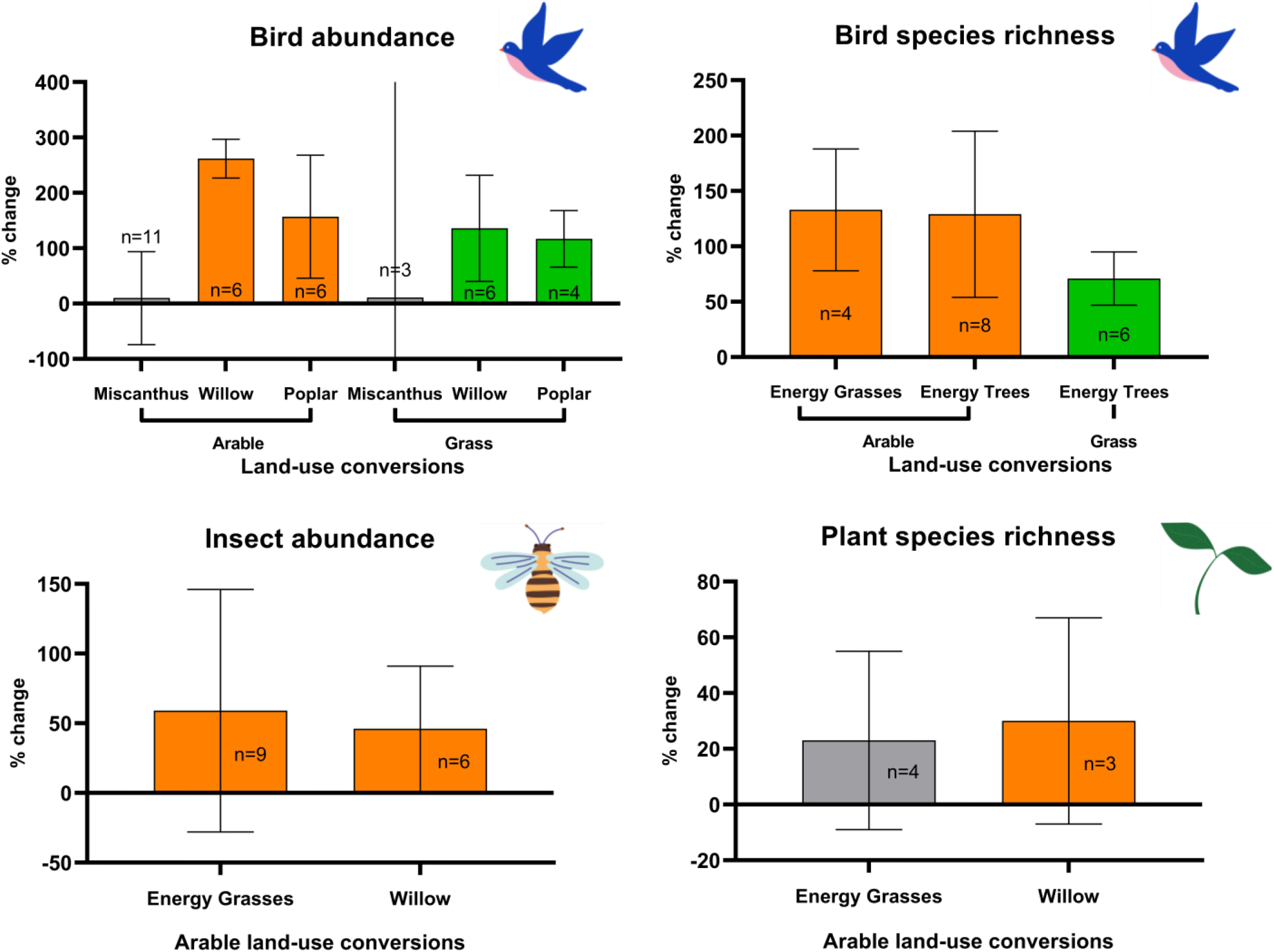
The impact on biodiversity species groups of specific land-use changes from agricultural land-use (arable or grassland) to specific bioenergy crops of poplar, willow, and *Miscanthus*, or short-rotation energy trees (poplar or willow), or energy grasses (*Miscanthus*, switchgrass, and prairie grass). Bars represent mean values, with 95 % CI shown. Sample size is shown by ‘n’. Grey bar indicates non-significant result (shown by the Grass-*Miscanthus* (p=0.77) and Arable-*Miscanthus* (p=0.92) conversions for bird abundance and Arable-Energy Grasses (p=0.15) conversion for plant species abundance). The error bar for the Arable-*Miscanthus* conversion impact on bird abundance does not fit on the axis. Separate axis scales are used for each pane.

### Land-use change to bioenergy: impacts on landscape amenity

Of the 2,364 papers screened just 12 addressed the specific question of the amenity and recreation impacts of bioenergy crops. Findings from these studies are summarised for the public and landowning stakeholders, as well as considerations for policymakers, in Figure 4. Two of the final 12 papers evaluated the visual impact of bioenergy crops alongside other impacts of bioenergy: air pollution, road traffic, and power station appearance were considered greater concerns of bioenergy infrastructure deployment than the landscape impact of the bioenergy crops themselves^32,39^. This was supported by a separate UK study finding bioenergy infrastructure more controversial with the public than bioenergy crops^30^. However, all three studies identified some public concern regarding the landscape impact of bioenergy crops, with ‘loss of view’ and ‘conspicuousness in the landscape’ both frequently mentioned as a concern in a study employing focus groups^25^ (see Table S2 for details of the 12 studies analysed).

**Figure 4.**
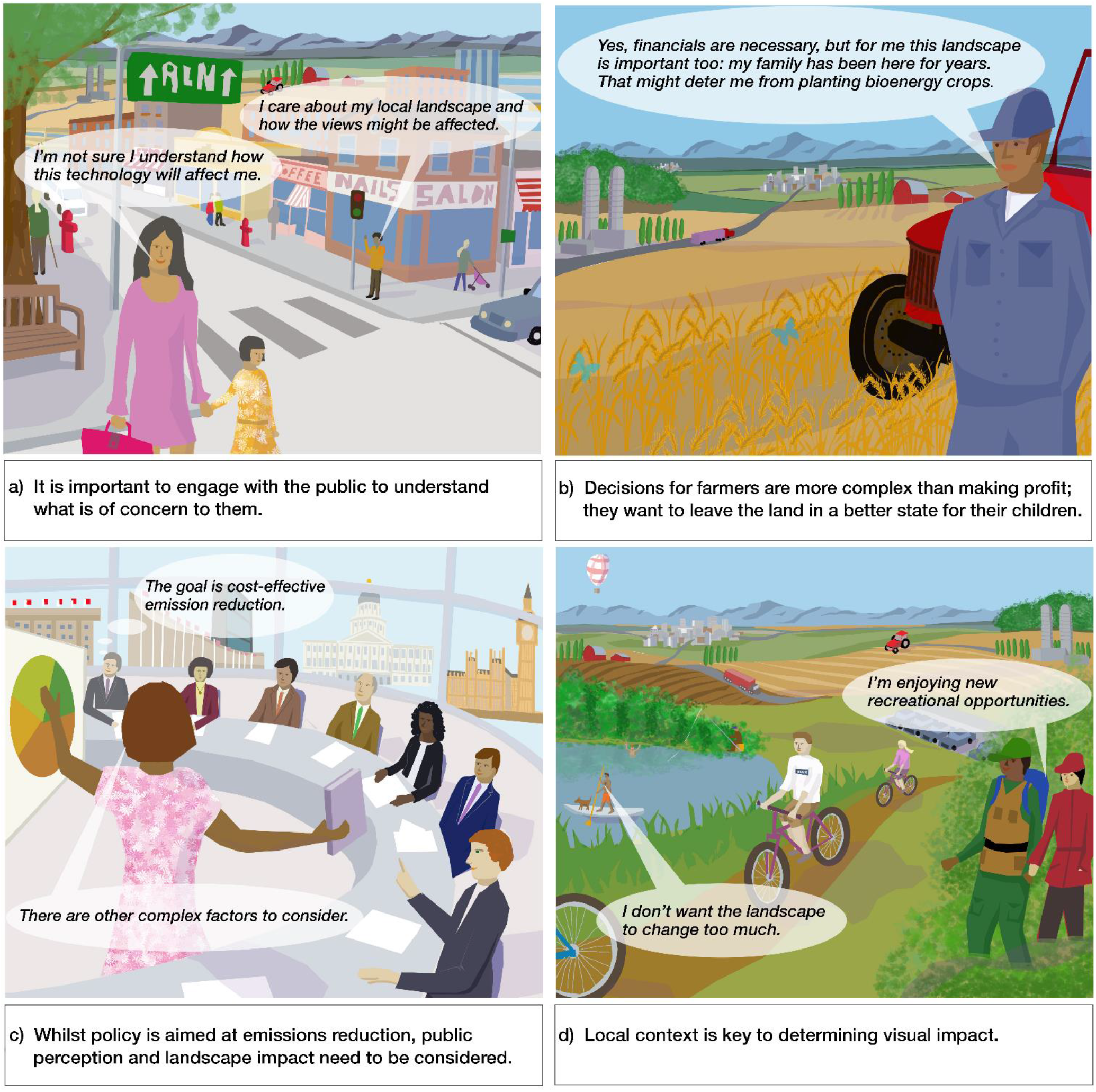
Landscape impacts of bioenergy crops are perceived differently by different stakeholders. Here we represent themes which emerged from the systematic review of amenity (visual and recreation) impact of planting bioenergy crops. a) reflects the lack of public understanding, engagement and information on the impact of bioenergy crops in local landscapes. b) summarises the values and motivations of farmers when considering whether to plant bioenergy crops on the landscape. c) local context is key to ladnscape decisions: who uses the landscape, how landscape features could change under bioenergy crop planting, and the diverse attitudes of people recreating in that landscape context. d) policymakers need to make decisions, such as meeting net-zero targets, whilst understanding the importance of other factors at the community level, which is challenging when these factors can be context specific and difficult to quantify.

Landscape context was found to shape attitudes towards visual impact of bioenergy crops^30,40–43^, with several reports showing that public attitudes to these new crops are contingent on the current landscape^40,41,43,44^. For example, Eaton *et al*.^40^ showed that bioenergy crops are supported in areas with existing trees, but may be opposed in open landscapes. In contrast, Dockerty *et al*.^30^ found evidence that visual amenity benefits from bioenergy crop deployment can be realised where deployment increased landscape complexity or heterogeneity, which could occur in more open landscapes. Such contrasting findings from different regions highlight the complexity of drawing overarching conclusions around impacts that may be highly context specific.

We found evidence in one large-scale survey that farmers who valued landscape amenity are less likely to be willing to plant bioenergy crops^45^. This contrasted with further evidence, from individual interviews, that landowners supported bioenergy crops because of visual attractiveness and the provision of hunting cover for wildlife^46^. These conflicting and limited results preclude a firm conclusion on farmer attitudes on the aesthetic impact of bioenergy crops. As with public attitudes, the role of study location and context are highly relevant.

It remains difficult to reach firm conclusions regarding the amentiy impact of large-scale deployment of bioenergy crops because people have often not been exposed to these land-use change. Some studies asked questions about potential bioenergy scenarios without the use of visual aids^32,39,40^, which raises the question of whether the individuals involved understood the nature of this landscape change. In contrast, using images^45, 29^ and 3D ‘real-time’ landscape models^29^ enabled interactive engagement for participant and since public awareness and deployment of bioenergy crops is relatively low^29^, such tools may have an important role in the future.

## Discussion

Future bioenergy policy will need public support to be successful and this will be shaped by public engagement, the nature of bioenergy expansion, and the impacts which follow^47,48^. In this first meta-analysis on the impacts on biodiversity of agricultural LUC to to bioenergy cropping, we have shown that increased deployment of these crops can deliver significant biodiversity benefits. This represents a key finding given the important role of biodiversity in underpinning critical ecosystem functions and services. It suggests that bioenergy cropping systems, that often utilize perennial rather annual crops, have the potential to be more stable and resilient than annual arable cropping, although long-term field studies are required to confirm this effect. In contrast, with little evidence on the relationship between bioenergy crops and landscape amenity, where public views are key, few conclusions could be drawn. This latter finding raises important questions about the acceptability of large-scale bioenergy crop planting for the public: for example, BECCS features heavily in the UK government’s approach to reaching a net-zero economy by 2050^49^, but the recent UK Climate Assembly found limited public support for BECCS^50^, showing the uphill challenge in achieving social legitimacy for this new and unproven technology. Our work highights a substantial knowledge gap in understanding when the public is most supportive of opportunities for deployment of the technology and associated bioenergy crops: it will be critical to close this gap to increase public acceptance.

### Biodiversity

Conversion of agricultural land to bioenergy crops results in improved biodiversity across species groups (Figure 2), providing novel insight into an important driver for ecosystem services and adding to the existing literature of other ecosystem services supported by bioenergy cropping, including soil organic carbon and flood mitigation^8,9,15,51,52^. Crucially, biodiversity could be supported alongside economic returns in a diversified farm business, particularly if future policy incentivises delivery of ecosystem services^53^. Future improved crop yields and dietary shifts away from animal agriculture could free up agricultural land for conversion to bioenergy crops, supporting biodiversity without undermining food security and could be an attractive way to expand bioenergy cropping, given that a recent review found evidence that LUC from natural ecosystems to bioenergy crops could be negative for biodiversity^19^.

Our findings point to trade-offs between biodiversity and food production: whilst converting arable land to bioenergy crops delivered greater biodiversity benefits than converting grassland, it also reflects a higher food production opportunity cost. Trade-offs may also exist at the level of bioenergy crop choice: conversions to energy trees (poplar and willow) delivered greater biodiversity benefits than conversions to energy grasses (*Miscanthus*, switchgrass, and prairie grass; Figure 3), although these results were driven by the large number of bird studies reviewed and there could be other reasons, such as yield or water-use considerations, to favour energy grasses in specific contexts. Whilst our results show the impact of specific LUC on biodiversity, these findings are generic and the exact impact on biodiveristy will also be determined by land management decisions, such as agrochemical inputs, weed control, tilling, vegetation structure, harvesting and crop rotation^19,54,55^.

One important caveat is that of scale, since most of our data were drawn from field studies generally under 10 ha, where bioenergy fields added to landscape hetroegeneity which is known to be important to supporting biodiversity^56,57^. Thus, the scaleability of these results is questionable, particularly when considering large monocultures of extensive bioenergy planting over several hundred hectares, where positive impacts on biodiversity could easily be reversed^59,60^. Understanding how these positive impacts on biodiversity can be preserved at landscale-scale remains the significant, yet solvable, challenge.

Our research here has only considered system-bound direct land use change to bioenergy cropping, rather than the indirect Land Use Change (iLUC): impacts that can occur if bioenergy crops displace food crops elsewhere, for example. These indirect or consequential impacts are beginning to be quantified in Consequential Life Cycle Analyses (C-LCA)^58^ but are outside the scope of this study.

### Amenity Value

Disparate research methods in the 12 amenity studies analyzed made it difficult to reach generic conclusions with respect to the impact of bioenergy cropping on amenity value. Only two of the studies involved rigorous public engagement^30,40^ and they each employed different approaches concerning their questions and use of visual aids. Although both found positive results regarding public attitudes towards the landscape impact of bioenergy crops, these were context specific and found alongside concerns on visual impact.

Such inconsistencies have been found in other assessments of landscape impact for different renewable energy technologies. A meta-analysis of public responses to wind energy turbines found a wide range of research methods and only limited agreement on a set of basic visual impact variables, preventing definitive conclusions^59^. A further wind energy study found evidence of public sensitivity to wind turbine placement in landscapes of high aesthetic value, and high acceptance in unattractive landscapes^29^, which could be used to inform bioenergy planting decisions. A review of previous studies to connect landscape features to aesthetic value concluded that no results have been achieved which could be translatable into policy. One of these studies, a meta-analysis of photograph-based perception found that no variable of landscape character had a clear relationship with aesthetic value^60^. Landscape heterogeneity positively affects aesthetic value, according to one meta-analysis, echoing a finding of our review, although their result was based on just six studies, and ours just one^30^. Similarly, another review of studies on human perceptions of the landscape provided a summary of landscape features supporting landscape aesthetics, supported by low numbers of studies^61^. Our findings reflect a challenge with the wider landscape amenity literature: results are not translatable out of the study context, and few studies seek to replicate previous work^62^. More recently, research has made use of geo-referenced social media data as a proxy for recreation visits to determine drivers of landscape amenity^63^. Echoing the above findings, key factors are typically not connected to landscape features, with population density, accessibility, proximity to water, and mountainous terrain all driving visitation rates^64^.

Policy guidelines on landscape amenity typically draw on expert opinion, not primary research^62^, and this is problematic because these guidelines may overlook aesthetic impacts^65^, as well as the potentially weak correlation between public views and expert views of determinants of landscape aesthetics^61^. Moving forward, policy could be guided by two paths. Firstly, avoiding deploying bioenergy crops in landscapes of high amenity value, where potential for negative impact appears greatest^66^. Secondly, facilitating local-level decision making and information dissemination to ensure that community-level attitudes are voiced and bioenergy crops are deployed in a way deemed acceptable by those who will be affected. Local-level engagement is identified as crucial to achieving a Social License to Operate^67^ and has proved critical to the success or failure of new energy technologies in achieving social legitimacy^68^. Community level engagement may also further our understanding of how bioenergy crops impact the ‘sense of place’ – the nature of the connection that we hold to the landscape^45^. Although no evidence of this was found from our review the sense of place has influenced attitudes towards other renewable energy technologies^69,70^.

Similar to the biodiversity studies, the amenity papers reviewed did not address the impact of landscape-scale bioenergy monocultures. With limited evidence suggesting that landscape heterogeneity increases visual amenity value, it could be expected that monocultures, which reduce heterogeneity, would undermine landscape amenity^71^.

### Conclusions

Meeting net-zero targets through ambitious plans for accelerated bioenergy crop planting requires important decisions to be made in balancing negative emissions, alongside food production and the natural environment. In this first meta-analysis to address the impacts of bioenergy cropping on biodiversity in agricultural landscapes, we find significant biodiversity improvements at the farm-scale across a number of diverse taxa, where land is converted from either arable or managed grassland to tree and grass bioenergy crops. In contrast, the amenity impacts of bioenergy cropping are harder to quantify with both positive and negative responses, depending on the specific context, reflecting a major knowledge gap. Taken together, these findings suggest that the proposed large-scale bioenergy and BECCS deployment under energy scenarios consistent with net-zero policies and the Paris targets could be compatible with improved farm-scale biodiversity but further research is required on scalability of results, approaches to landscape heterogeneity, alongside public acceptance and social legitimacy if these targets are to be met.

## Acknowledgements

We extend our thanks to the authors of the studies which were used in this analysis, and particularly to those authors who were reached for support and who were able to provide additional data and clarifications. This work was supported by the NERC-funded UK Energy Research Centre, by the NERC project Addressing the Valuation of Energy and Nature Together (ADVENT, NE/ M019764/1) and by The University of California, Davis with CD the recipient of a NERC PhD studentship (1790094). ZMH was supported by a NERC Industrial Innovation Fellowship (NE/R013314/1).

## Data Availability

We provide in the SI the data table of the final 12 studies used in our amenity review. Upon request of the authors an Excel table is available providing the full data-set used in the biodiversity meta-analysis.

