## Supplementary Information for "Land-use change from food to energy: meta-analysis unravels effects of bioenergy on biodiversity and amenity"

Supplementary Table 1 *Search strings used in Web of Science and Scopus as part of the two systematic reviews conducted in July 2020.*

| | |
|---|---|
| **Biodiversity search string:** | (salix OR miscanthus OR poplar OR willow OR populus OR switchgrass OR "phalaris arundinacea" OR "reed canary" OR "short rotation" OR SRC OR "woody biomass" OR "bioenergy crop*" OR "biomass crop*" OR "*energy crop*") |
| AND | ("land-use change" OR "land use change" OR bioenergy OR biomass OR biofuel OR BECCS OR CCS OR "carbon capture" OR "land-use change*") |
| AND | (biodiversity OR "species abundance" OR "species richness" OR "species distribution" OR ecology OR *fauna OR flora OR marine OR freshwater OR fresh-water OR nesting OR beetle* OR bird* OR mammal* OR microb* OR fung*) |

| | |
|---|---|
| **Amenity search string:** | (bioenergy OR biofuel OR "energy crop*" OR "woody crop*" OR "energy plantation*" OR "short-rotation" OR "woody biomass" OR "energy forest*" OR SRC OR willow OR Miscanthus OR poplar OR salix) |
| AND | (view OR visual OR visib* OR landscape OR recreation OR visit OR "public access" OR walk OR hunting) |
| AND | (valu* OR benefit OR amenit* OR alien OR invasion OR naturalness OR public OR "ecosystem service*" OR "environmental service*" OR "natural capital" OR attitude OR preference OR perception OR questionnaire OR interview) |

Supplementary Table 2 *Summary of the 12 papers analysed in the amenity systematic review*

| Study name | Study country | Bioenergy crop studied | Visual or recreational impact | Research method | Research question | Details of research methods | Main finding(s) |
|---|---|---|---|---|---|---|---|
| Upham and Shackley (2007) | UK | Miscanthus and willow | Visual (landscape) | Survey | Is landscape change to energy crops a concern for local people? | A total 573 questionnaires were completed by local residents in 2004. | Landscape impact is a public concern, but it is a lower priority concern than other concerns associated with new energy infrastructure. |
| Boll et al (2014) | Germany | SRC | Visual (landscape) | Interview | How do local people respond to landscape changes in their recreation areas, including introduction of bioenergy crops? | Interviews were held with 400 residents (10-15 minutes each); questions focussed on how a specific local recreation area would be affected by bioenergy crops. | The context of the existing landscape was key in determining whether people approved of bioenergy crops being grown, and whether it improved 'recreational suitability'. |
| Upham (2009) | UK | Miscanthus and willow | Visual (landscape) | Survey | Is landscape change to energy crops a concern for local people? | A total 290 questionnaires were completed by local residents in 2007, with very similar questions to the 2004 questionnaire in Upham and Shackley (2007). | Finds a rise in public concern for landscape impact of energy crops, but it remains of lower priority to other concerns associated with energy infrastructure. |
| Skarback and Becht (2005) | Sweden | Poplar | Visual (landscape) | Expert assessment | How does the introduction of energy forests affected the landscape and how it is perceived by people? | landscape analysis, using: field studies, maps, viewing points, line of sight, exposed landscape elements; photographic documentation: 35 photo points were recorded at each of the four seasons, over three years. | Viewing points (n=25) were assessed from the bioenergy crop: they were mostly not obstructed. |
| Dockerty et al (2012) | UK | SRC and Miscanthus | Visual (landscape) | Questionnaires, focus groups, interviews | What are public attitudes, particularly regarding visual impact, towards growing bioenergy crops in the landscape? | Photographs were used in questionnaires with local residents (East Midlands and South West England; 490 responses), with computer-generated visualisations of bioenergy landscapes used in focus groups (with 55 people). | Whilst some public conerns for impact of energy crops most respondents believed they could fit in well into a landscape, and power station infrastructure was of a higher concern. |
| Hipple and Duffy (2002) | USA | Switchgrass | Visual and Recreation | Interview | What motivates farmers to grow bioenergy crops? | A total of 52 farmers (existing and potential growers of switchgrass) were interviewed. | Farmers liked switchgrass for improved hunting cover, to attract more wildlife to their farm, and for its aesthetic qualities. |
| Bell and McIntosh (2001) | UK | SRC | Visual (landscape) and Recreational (footpaths) | Expert assessment | How does SRC affect the landscape character and how can it be structured so as to fit in with the landscape? | Recommendations of planting in different landscapes based on a review of 'practices on the ground'. | Context of the existing landscape is key in determining how and if bioenergy crops fit; variety in age and structure help large-scale planting fit into landscape. Footpaths can become more interesting. |
| Fawcett and Fawcett (2000) | UK | SRC | Visual (landscape) | Expert assessment | What are the landscape impacts of growing SRC and are existing guidelines on this appropriate? | Detailed site visits at different times of the year to eight case study sites of varying landscape type and sensitivity across England. Five further sites assessed on reconnaissance basis, all in Yorkshire. | Context is important in determining how bioenergy fits into landscape; nine of 14 diverse sites assessed found to be suitable for planting; scale and concentration of planting drive adverse affects. |
| Eaton et al (2018) | USA | Miscanthus, switchgrass, willow | Visual (community) and Recreation (wildlife viewing) | Survey | What are farmers' motivationsand values and how do these relate to willingness to grow bioenergy crops? | A total of 907 survey responses from farmers and landowners. Survey questions used the Likertscale (answers allowed four levels of agreement/disagreement). | Farmer concern for the impact of bioenergy crops on local community, those valuing recreation goals of land-use (view and wildlife habitat) were less likely to support planting these crops. |
| Sikorska et al (2020) | Poland | Miscanthus | Visual impact (urban park) | Interview | How do park users respond to the aesthetic value of their park replacing existing vegetation with energy crops? | Computerized visual simulations used to show park users how the park would look under bioenergy crops. | In an urban park environment the public preferred energy crops in place of grass; energy crops reduced noise pollution, and obscured views of the city. The energy crops appeared to add complexity to the park. |
| Sherrington et al (2008) | UK | SRC willow, Miscanthus | Visual impact (community) | Focus group | What are the barriers to farmers growing bioenergy crops? | A total of 31 farmers (existing and potential Miscanthus and SRC willow growers) were involved in three focus groups at locations across the UK, near to biomass power stations. | Farmers growing these bioenergy crops did not report any negative feedback from the public thus far. |
| Bell (1994) | UK | SRC | Visual impact (landscape) | Expert assessment | How do energy forests impact the landscape and how can they be structured so as to maintain aesthetic value of the landscape? | Application of the landscape architecture literature to determine optimal strategies for integrating bioenergy crops into different landscapes. | The context of the landscape determines how the energy crop should be structured so as to maintain landscape aesthetics. |

### Meta-analysis procedures

A number of judgement calls were required in order to deem whether and which data was includable in the meta-analysis. There issues were discussed between authors and are documented below, detailing the nature of the issue and the decision which was made.

Supplementary Table 3 *Details of judgement calls made during course of meta-analysis*

| **Paper affected** | **Issue** | **Decision** |
|---|---|---|
| Helms et al. 2020; Bellamy et al. 2009; Rowe et al. 2006 | A number of studies did not report on all insects found, but a sub-set. For Helms et al. (2019) only 'ant activity' data was given for insect abundance. Bellamy et al. (2009) reported 'all invertebrates', Rowe et al. (2006) reported 'predatory arthropods'. | These data were used in the insect abundance meta-analysis. |
| Lagerlof et al. 2014 | Formalin was used to count earthworm abundance in several studies. | Any data derived from formalin was discounted because this is generally considered an unreliable way to count earthworms, and opposed to the standard practice of hand counting. |
| Sage et al. 2010 | A Standard Error or mean value of '0' was reported for several data points in the results but a zero value is not usable in meta-analysis. | Standard protocol was followed here, with a constant '0.5' added to all values (mean and error values, of treatment and control) for this study where zero values occurred. |
| Verheyen et al. 2014 | 'Activity density' was used instead of 'insect abundance'. | These data were used for insect abundance, with the two terms deemed interchangeable. |
| Christian et al. 1997 | Christian et al. 1997 calculated 'expected' bird species richness using rarefaction. This methodology can be problematic because bird species richness is likely to be different at | Notwithstanding the issues with using rarefaction we decided to use the data in the meta-analysis, and the data were not an outlier in the results. |

| | | |
|---|---|---|
| | different scales and modelling this using a rarefaction curve could give inaccurate results. | |
| Werling et al. 2014; Gardiner et al. 2010; Helms et al. 2020; Orr et al. 2015; Liang et al. 2012; Oates et al. 2016; Jesus et al. 2016. | We were primarily interested in energy crops of poplar, willow, switchgrass, and *Miscanthus*. However, our systematic search returned a number of papers which also looked at the potential bioenergy crop prairie grass. | We determined to use the data reported in these papers, as they were returned by our systematic search, and represent an energy crop comparable to *Miscanthus* and switchgrass. |
| Chiatante et al. 2019 | 'SRCcut' is reported as well as 'SRC'. | Data were used for both these values as a cut SRC field represents a temporal state in which the SRC field will be in during its commercial development for bioenergy. Values were not reported in other studies for the cut period of the energy tree field. |
| Verheyen et al. 2014 | Nine of the ten treatment sites were poplar sites, and one of the ten sites was a willow site. | The treatment was listed as poplar, given this was the dominant crop used in the treatment. |
| Feledyn-Szewczyk et al. 2019 | Reporting of n value is important for meta-analysis and most papers were clear on this. We disagreed with how Feledyn-Szewczyk et al. reported n for their study, and it did not align with how other studies reported n (typically the number of fields, or field sites). Feledyn-Szewczyk et al. instead included multiple data points from the same location, treating these points as independent suggesting the potential for pseudoreplication. They reported n=30, when we deemed n=5 appropriate, corresponding to the number of field plots. | We used n as reported by Feledyn-Szewczyk et al. because whilst we disagreed with this, their n-value will have influenced other statistics reported in their paper which we went on to use. And so for consistency we used n=30. However, we did check to see the impact that this had on our results, with the effect minimal. |
| Chiatante et al. 2019 | Chiatante et al. reported n as the number of point counts within field plots, and not the number of fields in their study. We considered this for the potential of pseudoreplication. | We decided to use n as reported by Chiatante et al. because the point counts were taken from large field sites, and because the n-value used in the paper will have influenced other statistics that we used in our meta-analysis. |
| Bellamy et al. 2009 | Abundance was reported for 'adult earthworms' and not 'earthworms'. | These data were used for the earthworm abundance meta-analysis. |
| McGowan et al. 2019; Orr et al. 2015; Jesus et al. 2016; Oates et al. 2016; Liang et al. 2012. | Soil microbial biomass was reported using microbial phospholipid fatty acid (PLFA) profiles. This was reported slightly differently between studies: McGowan et al. 2019 reported 'total PLFA'; Orr et al. 2015 reported 'total PLFA-FAME abundance' Jesus et al. 2016, Oates et al. 2016, and Liang et al. 2012 reported 'total microbial lipids'. | All these means of assessing soil microbial biomass were deemed comparable for the meta-analysis. |

### Study heterogeneity calculations

A heterogeneity test was used to check the studies used in our meta-analysis were sufficiently comparable in study-design to allow comparison across them using meta-analysis. Cochran's Q test was used to determine that the data was sufficiently heterogeneous. We also looked at the $I^2$ value: $I^2$ is the percentage of variation across studies which is due to heterogeneity not chance, with a higher $I^2$ suggesting higher heterogeneity. The meta-analysis software OpenMEE was used to calculate the Q and $I^2$ statistics.

### Publication bias – Funnel plots

Funnel plots are given for each of the meta-analyses carried out below. The absence of publication bias is identified by a symmetrical funnel with a larger spread for smaller sample sizes.

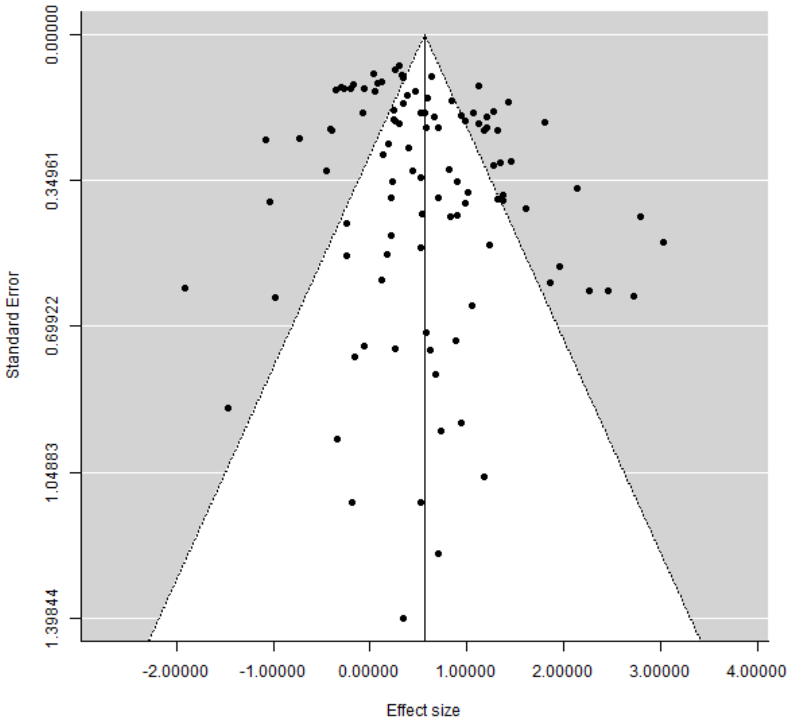


Supplementary Figure 1 *All data points funnel plot*

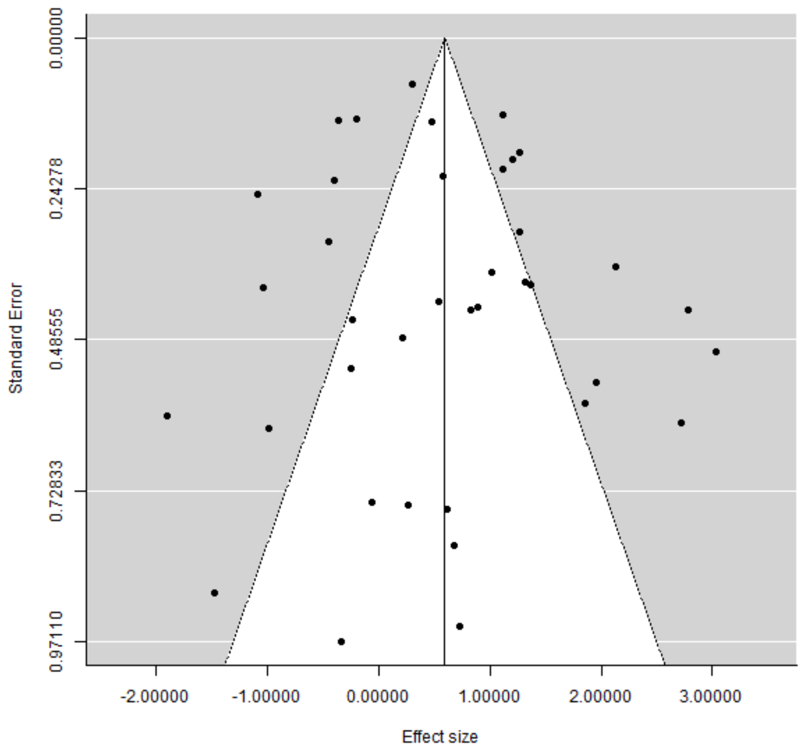


Supplementary Figure 2 *Bird abundance funnel plot*

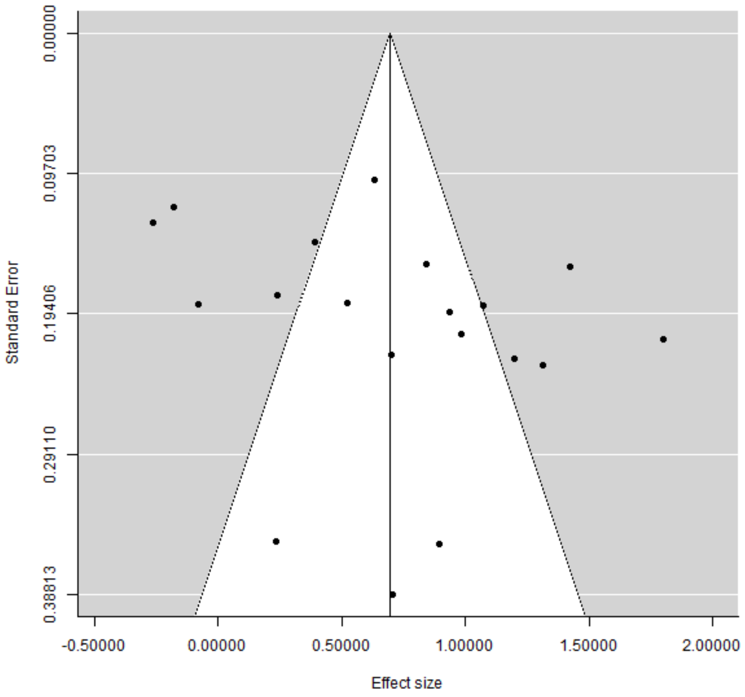


Supplementary Figure 3 *Bird species richness funnel plot*

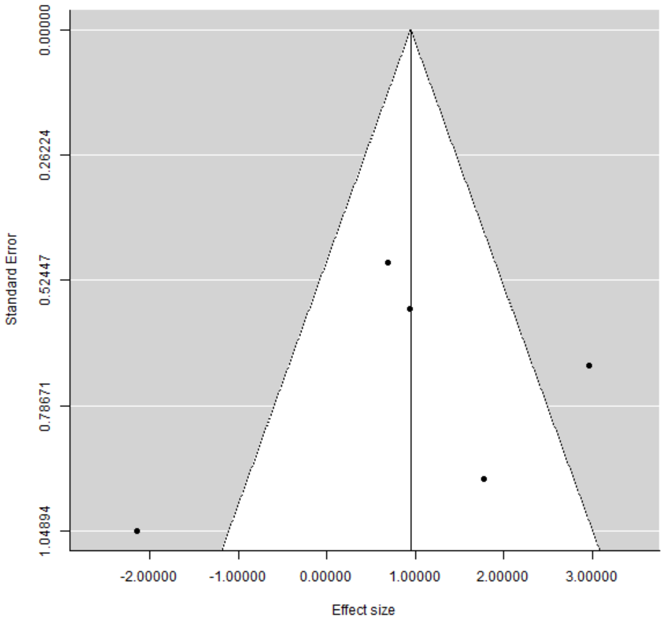


Supplementary Figure 4 *Buntings abundance funnel plot*

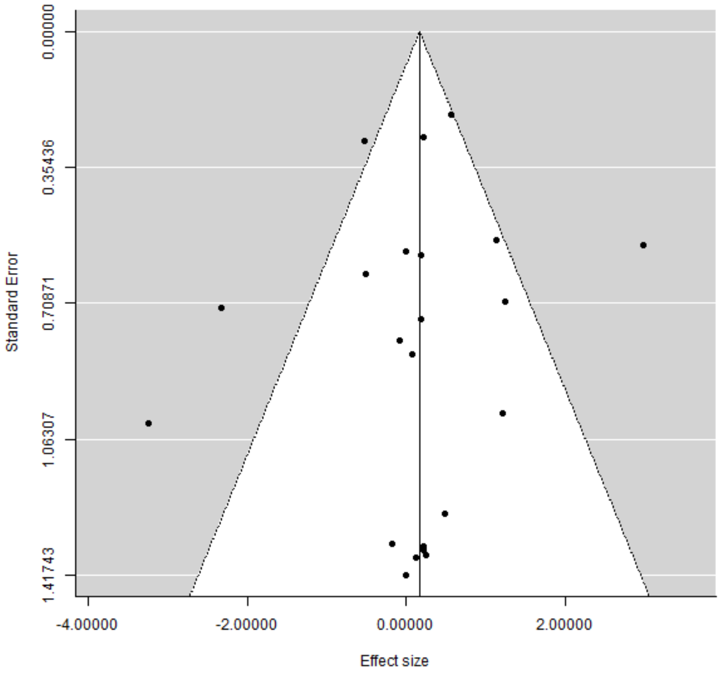


Supplementary Figure 5 *Skylark abundance funnel plot*

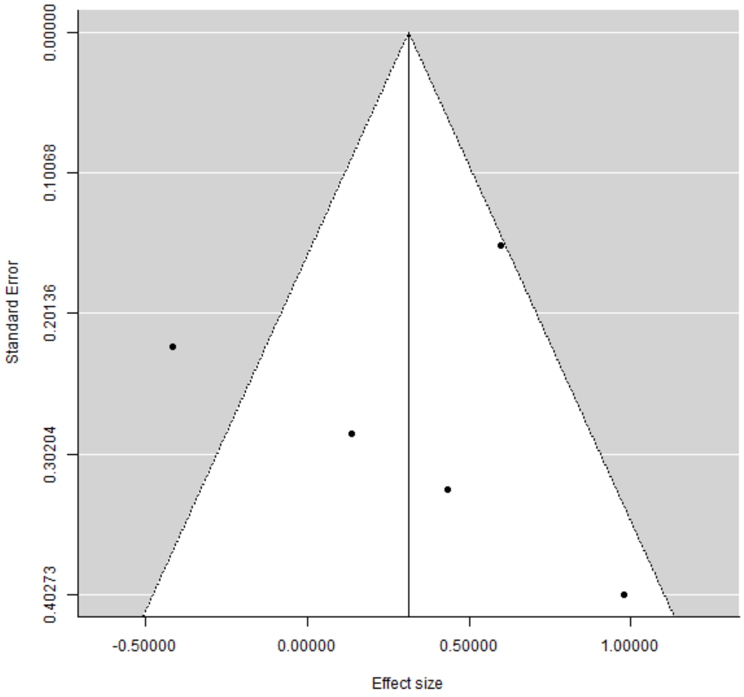


Supplementary Figure 6 *Earthworm abundance funnel plot*

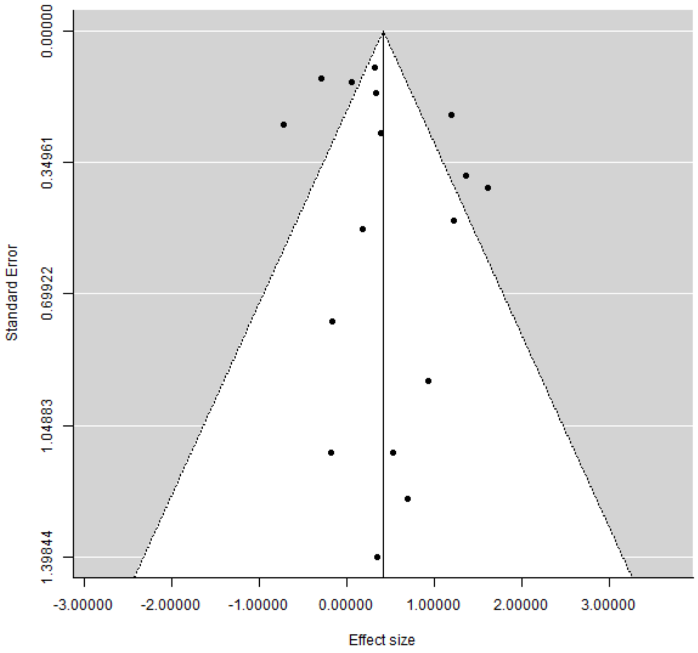


Supplementary Figure 7 *Arthropod abundance funnel plot*

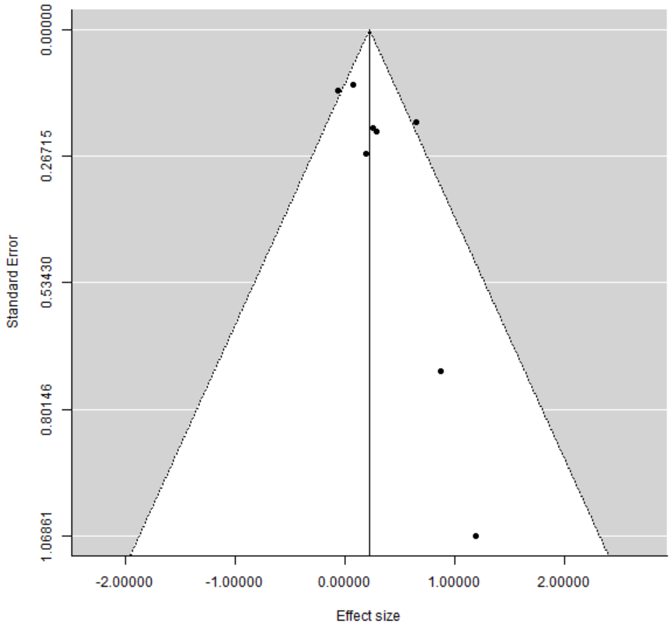


Supplementary Figure 8 *Plant species richness funnel plot*

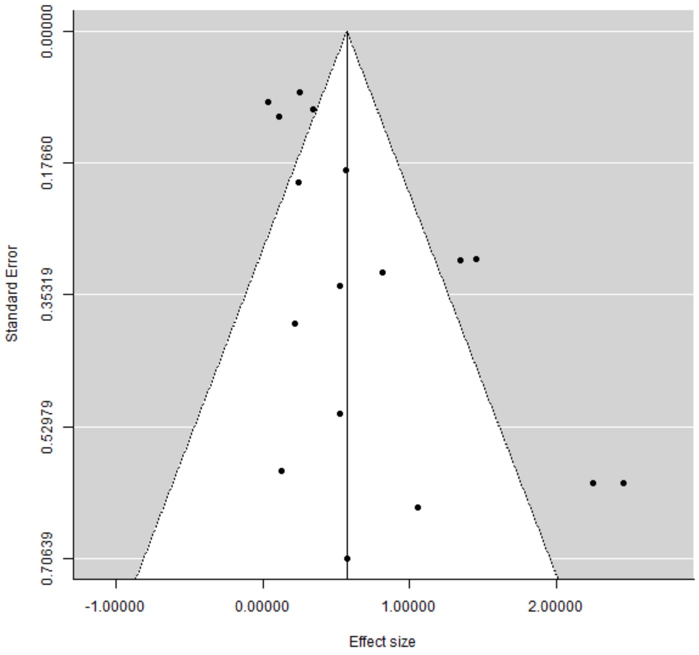


Supplementary Figure 9 *Soil microbial biomass funnel plot*

**Meta-analysis calculations**

The OpenMEE software was used to calculate the log ratio of biodiversity between the agricultural land-use and the bioenergy cropping land-use, and the standard error:

$$\ln R = \ln(\text{mean}(\text{bioenergy})) - \ln(\text{mean}(\text{control}))$$

$$SE = (CI(\text{high}) - \ln R)/1.96$$

Where *lnR* is the log ratio, *SE* is the standard error, and *CI* is the confidence interval.

This effect size and its standard error were then converted into a percentage change for ease of demonstration of changes between the control and treatment groups:

$$\ln R(\%) = \text{Exp}(\ln R) - 1$$

$$SE(\%) = \text{Exp}(SE) - 1$$

We calculated the percentage change confidence intervals according to the following equation:

$$CI\%(\text{high}) = \text{Ln}R(\%) + SE(\%)$$

$$CI\%(\text{low}) = \text{Ln}R(\%) - SE(\%)$$

**Supplementary Meta-analysis results**

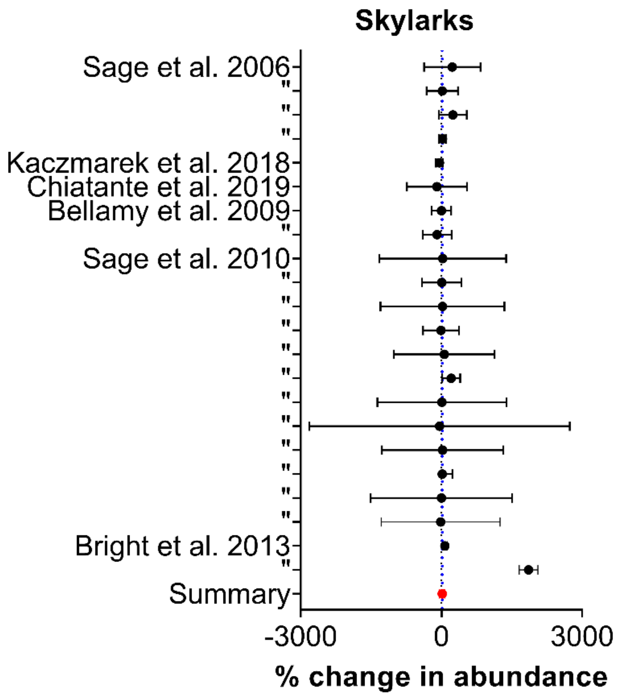


Supplementary Figure 9 *Forest plot of skylark meta-analysis (n=22). Summary effect = 18 % ± 61 %, p=0.50.*

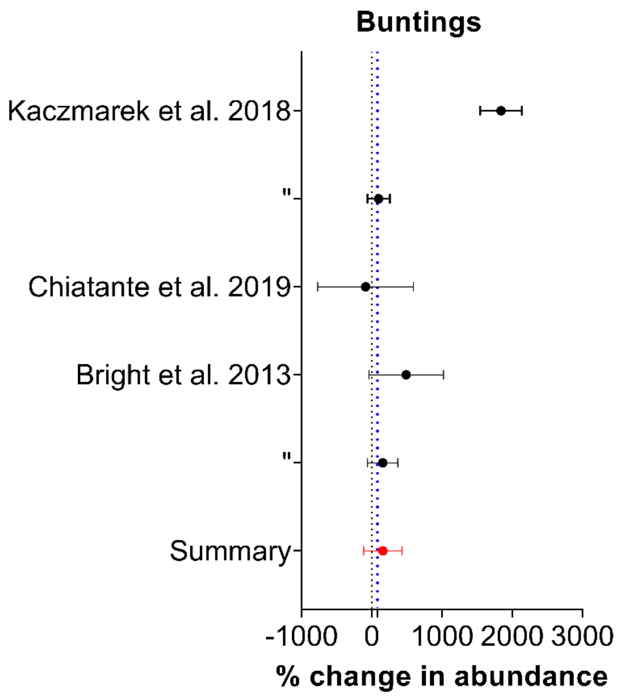


Supplementary Figure 10 *Forest plot of bunting meta-analysis (n=5). Summary effect = 158 % ± 271 %, p=0.16.*

### Data limitations

An important knowledge gap concerns the bioenergy cropping impact on mammals, with very limited understanding of the impact of bioenergy land-use change upon mammals, as noted by previous research (Riffell et al., 2011; Rowe et al., 2009). A further limitation of our results is the heavy concentration of studies in the USA and UK, with a focus on specific bioenergy crops (*Miscanthus*, switchgrass, poplar, and willow) and the uncertainty of whether these results would be reflected in other regions and with other bioenergy crops. Further studies in other countries and regions would help develop understanding of the bioenergy crop impact on biodiversity.

### Studies used in the biodiversity meta-analysis and amenity systematic review


Bell, S. (1994). Energy forest cultivation and the landscape. *Biomass and Bioenergy*, *6*(1–2), 53–61. https://doi.org/10.1016/0961-9534(94)90085-X

Bell, S., & McIntosh, E. (2001). Short Rotation Coppice in the Landscape, 1–8.

Bellamy, P. E., Croxton, P. J., Heard, M. S., Hinsley, S. A., Hulmes, L., Hulmes, S., … Rothery, P. (2009). The impact of growing miscanthus for biomass on farmland bird populations. *Biomass and Bioenergy*, *33*(2), 191–199. https://doi.org/10.1016/j.biombioe.2008.07.001

Berkley, N. A. J., Hanley, M. E., Boden, R., Owen, R. S., Holmes, J. H., Critchley, R. D., … Parmesan, C. (2018). Influence of bioenergy crops on pollinator activity varies with crop type and distance. *GCB Bioenergy*, *10*(12), 960–971. https://doi.org/10.1111/gcbb.12565

Boll, T., von Haaren, C., & Albert, C. (2014). How do urban dwellers react to potential landscape changes in recreation areas? A case study with particular focus on the introduction of dendromass in the Hamburg Metropolitan Region. *IForest*, *7*(7), 423–433. https://doi.org/10.3832/ifor1173-007

Bright, J. A., Anderson, G. Q. A., Mcarthur, T., Sage, R., Grice, P. V, Bradbury, R. B., … Sage, R. (2013). Bird use of establishment-stage Miscanthus biomass crops during the breeding season in England Bird use of establishment-stage Miscanthus biomass crops during the breeding season in England, *3657*. https://doi.org/10.1080/00063657.2013.790876

Chiatante, G., & M. G., & Meriggi, A. (2019). Bird Diversity in Short Rotation Coppice in Northern Italy. *Ardea*, *107*(1), 5. https://doi.org/10.5253/arde.v107i1.a10

Christian, D. P., Collins, P. T., Hanowski, J. M., & Niemi, G. J. (1997). Bird and Small Mammal

Use of Short-Rotation Hybrid Poplar Plantations Author ( s ): Donald P . Christian , Patrick T . Collins , Joann M . Hanowski and Gerald J . Niemi Published by : Wiley on behalf of the Wildlife Society Stable URL : http://www.jstor.o, *61*(1), 171–182. Retrieved from https://doi.org/10.2307/3802426

Dockerty, T., Appleton, K., & Lovett, A. (2012). Public opinion on energy crops in the landscape : Considerations for the expansion of renewable energy from biomass. *Journal of Environmental Planning and Management*, *55*(9). https://doi.org/10.1080/09640568.2011.636966

Eaton, W. M., Burnham, M., Hinrichs, C. C., Selfa, T., & Yang, S. (2018). How do sociocultural factors shape rural landowner responses to the prospect of perennial bioenergy crops? *Landscape and Urban Planning*, *175*(August 2017), 195–204. https://doi.org/10.1016/j.landurbplan.2018.02.013

Emmerling, C. (2014). Impact of land-use change towards perennial energy crops on earthworm population. *Applied Soil Ecology*, *84*(March), 12–15. https://doi.org/10.1016/j.apsoil.2014.06.006

Fawcett, & Fawcett. (2005). *Assessment of the visual impacts of SRC plantations in Summaries of Biomass Projects carried out as part of the DTI's Technology Programme: New and Renewable Energy (eds: Duffy, G, & Beale, N.). DTI* (Vol. 176). https://doi.org/10.1140/epjst/e2009-01152-1

Feledyn-szewczyk, B., Matyka, M., & Staniak, M. (2019). Comparison of the E ff ect of Perennial Energy Crops and Agricultural Crops on Weed Flora Diversity.

Gardiner, M. A., Tuell, J. K., Isaacs, R., Gibbs, J., Ascher, J. S., & Landis, D. A. (2010). Implications of three biofuel crops for beneficial arthropods in agricultural landscapes. *Bioenergy Research*, *3*(1), 6–19. https://doi.org/10.1007/s12155-009-9065-7

Helms, J. A., Ijelu, S. E., Wills, B. D., Landis, D. A., & Haddad, N. M. (2020). Ant biodiversity and ecosystem services in bioenergy landscapes. *Agriculture, Ecosystems and Environment*, *290*(December), 106780. https://doi.org/10.1016/j.agee.2019.106780

Hipple, P. C., & Duffy, M. D. (2002). Farmers' Motivations for Adoption of Switchgrass. *Trends in New Crops and New Uses*, (1), 252–266.

Kaczmarek, J. M., Mizera, T., & Tryjanowski, P. (2019). Energy crops affecting farmland birds in Central Europe: insights from a miscanthus-dominated landscape. *Biologia*, *74*(1), 35–44. https://doi.org/10.2478/s11756-018-0143-1

Lagerlöf, J., Pålsson, O., & Arvidsson, J. (2014). Acta Agriculturae Scandinavica , Section B - Soil & Plant Science Earthworms influenced by reduced tillage , conventional tillage and energy forest in Swedish agricultural field experiments, (January 2011). https://doi.org/10.1080/09064710.2011.602717

Liang, C., Jesus, E. da C., Duncan, D. S., Jackson, R. D., Tiedje, J. M., & Balser, T. C. (2012). Soil microbial communities under model biofuel cropping systems in southern Wisconsin, USA: Impact of crop species and soil properties. *Applied Soil Ecology*, *54*, 24–31. https://doi.org/10.1016/j.apsoil.2011.11.015

McGowan, A. R., Nicoloso, R. S., Diop, H. E., Roozeboom, K. L., & Rice, C. W. (2019). Soil organic carbon, aggregation, and microbial community structure in annual and perennial biofuel crops. *Agronomy Journal*, *111*(1), 128–142. https://doi.org/10.2134/agronj2018.04.0284

Oates, L. G., Duncan, D. S., Sanford, G. R., Liang, C., & Jackson, R. D. (2016). Bioenergy cropping systems that incorporate native grasses stimulate growth of plant-associated soil microbes in the absence of nitrogen fertilization. *Agriculture, Ecosystems and Environment, 233*, 396–403. https://doi.org/10.1016/j.agee.2016.09.008

Orr, M. J., Gray, M. B., Applegate, B., Volenec, J. J., Brouder, S. M., & Turco, R. F. (2015). Transition to second generation cellulosic biofuel production systems reveals limited negative impacts on the soil microbial community structure. *Applied Soil Ecology*, *95*, 62–72. https://doi.org/10.1016/j.apsoil.2015.06.002

Riffell, S., Verschuyl, J., Miller, D., & Wigley, T. B. (2011). A meta-analysis of bird and mammal response to short-rotation woody crops. *GCB Bioenergy*, *3*(4), 313–321. https://doi.org/10.1111/j.1757-1707.2010.01089.x

Rowe, R. L., Goulson, D., Doncaster, C. P., & Clarke, D. J. (2013). Evaluating ecosystem processes in willow short rotation coppice bioenergy plantations, 257–266. https://doi.org/10.1111/gcbb.12040

Rowe, R. L., Hanley, M. E., Goulson, D., Clarke, D. J., Doncaster, C. P., & Taylor, G. (2010). Potential benefits of commercial willow Short Rotation Coppice ( SRC ) for farm-scale plant and invertebrate communities in the agri-environment. *Biomass and Bioenergy*, *35*(1), 325–336. https://doi.org/10.1016/j.biombioe.2010.08.046

Rowe, R. L., Street, N. R., & Taylor, G. (2009). Identifying potential environmental impacts of large-scale deployment of dedicated bioenergy crops in the UK. *Renewable and Sustainable Energy Reviews*, *13*(1), 271–290. https://doi.org/10.1016/j.rser.2007.07.008

Sage, R., Cunningham, M., & Boatman, N. (2006). Birds in willow short-rotation coppice compared to other arable crops in central England and a review of bird census data from energy crops in the UK. *Ibis*, *148*(SUPPL. 1), 184–197. https://doi.org/10.1111/j.1474-919X.2006.00522.x

Sage, R., Cunningham, M., Haughton, A. J., Mallott, M. D., Bohan, D. A., Riche, A., & Karp, A. (2010). The environmental impacts of biomass crops: Use by birds of miscanthus in summer and winter in southwestern England. *Ibis*, *152*(3), 487–499. https://doi.org/10.1111/j.1474-919X.2010.01027.x

Sherrington, C., Bartley, J., & Moran, D. (2008). Farm-level constraints on the domestic supply of perennial energy crops in the UK. *Energy Policy*, *36*(7), 2504–2512. https://doi.org/10.1016/j.enpol.2008.03.004

Sikorska, D., Macegoniuk, S., Łaszkiewicz, E., & Sikorski, P. (2020). Energy crops in urban parks as a promising alternative to traditional lawns – Perceptions and a cost-benefit analysis. *Urban Forestry and Urban Greening*, *49*(December 2019). https://doi.org/10.1016/j.ufug.2019.126579

Skärbäck, E., & Becht, P. (2005). Landscape perspective on energy forests. *Biomass and Bioenergy*, *28*, 151–159. https://doi.org/10.1016/j.biombioe.2004.08.008

Upham, P. (2009). Applying environmental-behaviour concepts to renewable energy siting controversy: Reflections on a longitudinal bioenergy case study. *Energy Policy*, *37*(11), 4273–4283. https://doi.org/10.1016/j.enpol.2009.05.027

Upham, P., & Shackley, S. (2007). Local public opinion of a proposed 21.5 MW(e) biomass gasifier in Devon: Questionnaire survey results. *Biomass and Bioenergy*, *31*(6), 433–441. https://doi.org/10.1016/j.biombioe.2007.01.017

Verheyen, K., Buggenhout, M., Vangansbeke, P., De Dobbelaere, A., Verdonckt, P., & Bonte, D. (2014). Potential of short rotation coppice plantations to reinforce functional biodiversity in agricultural landscapes. *Biomass and Bioenergy*, *67*, 435–442. https://doi.org/10.1016/j.biombioe.2014.05.021

Werling, B. P., Dickson, T. L., Isaacs, R., Gaines, H., Gratton, C., Gross, K. L., … Landis, D. A. (2014). Perennial grasslands enhance biodiversity and multiple ecosystem services in bioenergy landscapes. *Proceedings of the National Academy of Sciences*, *111*(4), 1652–1657. https://doi.org/10.1073/pnas.1309492111
